# Structure of the yeast ceramide synthase

**DOI:** 10.1101/2023.10.24.563787

**Authors:** Jan-Hannes Schäfer, Lena Clausmeyer, Carolin Körner, Bianca M. Esch, Verena N. Wolf, Stefan Walter, Dovile Januliene, Arne Moeller, Florian Fröhlich

## Abstract

Ceramides play a pivotal role as essential lipids, serving as foundational components for complex sphingolipids and potent signaling molecules. Ceramides are the products of the N-acylation of a sphingoid base and a CoA-activated fatty acid. This reaction is catalyzed by the enzymes of the evolutionarily conserved ceramide synthase (CerS) family. Yet, the precise structural details and catalytic mechanisms of CerSs have remained elusive. Here, we employed cryo-EM single particle analysis to unravel the structure of the yeast ceramide synthase complex in both an active and a fumonisin B1 inhibited state. Our findings shed light on the complex’s architecture, revealing a dimer of Lip1 subunits bound to the two catalytic subunits, Lag1 and Lac1. Each catalytic subunit forms a hydrophobic crevice that is accessible from both the cytosolic site as well as from the intermembrane space of the endoplasmic reticulum (ER). Within this cavity, we identify amino acids forming the active center and a sphingoid base, one of the substrates of the complex. Together, this suggests a pre-loaded state of the CerS. Additionally, the fumonisin B1 bound structure reveals the inhibitory mechanism by blocking the cytosolic acyl-CoA binding site.

## Introduction

Ceramides are pivotal players in sphingolipid metabolism^1^. They are active signaling molecules intricately involved in vital cellular processes^2,3^. Moreover, they function as critical intermediaries in the biosynthesis of complex sphingolipids like sphingomyelin and glycosphingolipids^4^. The accumulation of ceramides in metabolic tissues, particularly in the liver and skeletal muscle, disrupts essential processes such as insulin signaling^5,6^, fatty acid metabolism^5,7^, and mitochondrial function^7^. Dysregulation of ceramide synthesis is associated with various diseases, including cancer^8–10^, neurodegenerative disorders^11^, and ichthyosis^12^.

Ceramide synthases (CerS), a class of membrane-embedded enzymes, produce ceramides from a sphingosine backbone and an acyl-CoA^13–15^. Sphingolipid biosynthesis begins in the endoplasmic reticulum (ER) with the condensation of serine and palmitoyl-CoA, resulting in the formation of a short-lived intermediate, 3-ketosphinganine (3-KS)^16^. The first and rate-limiting step of sphingolipid biogenesis is catalyzed by the serine palmitoyl transferase (SPT) in yeast consisting of the core subunits Lcb1 and Lcb2^17^. It forms the so-called SPOTS complex with its interaction partners Tsc3, Orm1/2, and Sac1^18,19^. 3-KS is directly reduced to dihydrosphingosine (DHS), which, in yeast, can be further hydroxylated to phytosphingosine (PHS). DHS and PHS are substrates for the CerS in yeast^20^. In mammalian cells, de novo synthesized DHS (or sphinganine) and sphingosine, generated in the recycling pathway, are substrates for the N-acylation reaction catalyzed by the CerSs. In both cases, ceramides are transported from the ER to the Golgi apparatus, where they are further modified to yield the complex sphingolipids^21–24^.

All CerSs are ER-resident transmembrane proteins. They carry a TRAM-Lag1p-CLN8 (TLC) domain with six transmembrane helices^25^. Mammalian cells express six different CerSs, CerS1-6, that are also tissue-specific in organisms. The six different CerSs have different acyl chain lengths optima^26–29^. CerS5 and CerS6 prefer C16:0-, CerS1 C18:0-, and CerS2 and CerS4 prefer both C22:0- and C24:0-acyl-CoA substrates. Only CerS3 exhibits a broad specificity. The fatty acyl-CoA specificity of the CerSs is thought to depend on a short 11-amino acid loop, but the molecular details of this selectivity remain largely unknown^30^. In addition to the TLC domain, CerSs 2-6 also express a short homeo-domain that might function as a transcriptional regulator^31^. Yeast cells express two catalytically active ceramide synthases, Lag1 and Lac1, harboring the typical TLC domain. Both Lag1 and Lac1 carry two additional N-terminal transmembrane helices but miss the homeo-domain of the mammalian CerSs. Lag1 and Lac1 seem redundant in their function as only deleting both genes is lethal for yeast cells^32^. In addition to the catalytically active CerSs, a small yeast-specific subunit, Lip1, is required for CerS activity^33^. Its gene *LIP1* is essential, and its product is a single membrane-spanning protein that interacts with Lag1 and Lac1 to form an active CerS complex, but the stoichiometry and mechanism of activation are unknown.

Various cues control CerSs activity, including phosphorylation. The yeast CerSs activity is upregulated by Ypk1-dependent phosphorylation^34^. The Ypk1 kinases are master regulators of sphingolipid metabolism. Furthermore, Lag1 and Lac1 are subject to phosphorylation by yeast casein kinase 2 (CK2), which is required for optimal CerS activity^35^. In the case of human CerS2-CerS6, phosphorylation occurs at their C-terminal regions. However, the specific molecular changes induced by phosphorylation remain unclear. Additionally, mammalian CerSs are regulated through dimerization, facilitated by a short, evolutionary conserved, C-terminal DxRSDxE motif^36^. Importantly, the molecular details of CerSs’ active center remain elusive. Notably, two adjacent histidine residues are conserved in the CerS family required for catalytic activity^25,37^. A recent computational study proposed the existence of large hydrophobic cavities within a predicted model of human CerS and suggested a model for lateral substrate delivery to the enzyme from the membrane^38^.

Some CerS inhibitors are available. A mycotoxin produced by *Fusarium moniliforme* and related fungi is Fumonisin B1 (FB1)^39^. Fumonisins are 1-deoxy-analogs of sphinganines that share structural similarities to the sphingoid bases. They are potent inhibitors of ceramide synthases and are not isoform-specific. However, the molecular mechanism of CerS inhibition remains unknown. Structural information on the complex is required to gain mechanistic insights into the catalytic mechanism of CerS-mediated N-acylation of sphingoid bases.

Here, we solved the structure of the yeast CerS complex using cryo-EM single particle analysis. The CerS complex is heterotetrametric with pseudo-C2 symmetry. It comprises the two catalytic subunits Lag1 and Lac1 and two Lip1 small subunits. Furthermore, we solved the fumonisin B1 bound inhibited conformation of the complex. The interface of both halves of the complex is formed by the Lip1 homodimer, in which the subunits interact exclusively through their ER-luminal soluble domains. Lac1 and Lag1 are associated to the single transmembrane helices of each respective Lip1 subunit. Importantly, this configuration prohibits direct contact between the catalytic Lac1 and Lag1 subunits postulated for human CerSs. Each catalytic subunit contains a large cavity formed by the transmembrane helices 3-8. Its central position renders it accessible from the cytosol and the inter-membrane space facing the Lip1 single membrane helix. The two H residues required for catalytic activity are positioned in the center of the cavity. In addition, our fumonisin B1 inhibited structure reveals the mechanism of inhibition of the complex. Here, we provide the first available structures of a CerS, which will help resolve the molecular mechanism of ceramide biosynthesis.

## Results

To determine the structure of the yeast CerS, we engineered a yeast strain expressing all three subunits, Lag1, Lac1, and Lip1, under the control of the *GAL1* promoter. Lip1 was N-terminally tagged with a 3xFLAG tag. We tagged the Lip1 subunit on its N-terminus since it’s the only subunit essential for CerS activity in yeast^33^. For cryo-EM studies, the CerS was solubilized in glyco-diosgenin (GDN) and purified by FLAG-based affinity chromatography. The purification led to a stable complex, and the presence of all three subunits was confirmed by mass spectrometry-based proteomics (Sup. Fig. 1C). The purified complex showed CerS activity *in vitro*, in the presence of its substrates PHS and C26-CoA and was sensitive to the CerS inhibitor fumonisin B1. The addition of the purified yeast acyl-CoA binding protein Acb1 further increased the activity of the complex (Sup. Fig. 1I).

### Architecture of the ceramide synthase complex

Single particle cryo-EM was used to reveal the architecture of the yeast CerS complex, which was refined to 3 Å resolution (Fig.1, Sup. Fig. 2, Sup. Tab. 1). Our structural analysis revealed a pseudo-C2-symmetric hetero-tetramer formed by two subunits of the single-pass transmembrane protein Lip1 and one of each respective catalytic subunits Lag1 and Lac1 in each half. The interface is exclusively established through the soluble ectodomain of both Lip1 subunits, leaving a large gap between the transmembrane domains of each half. The small soluble domains of Lip1 consist of four antiparallel β-sheets, which are sandwiched by two small α-helices (Fig. 1E).

**Fig. 1:**
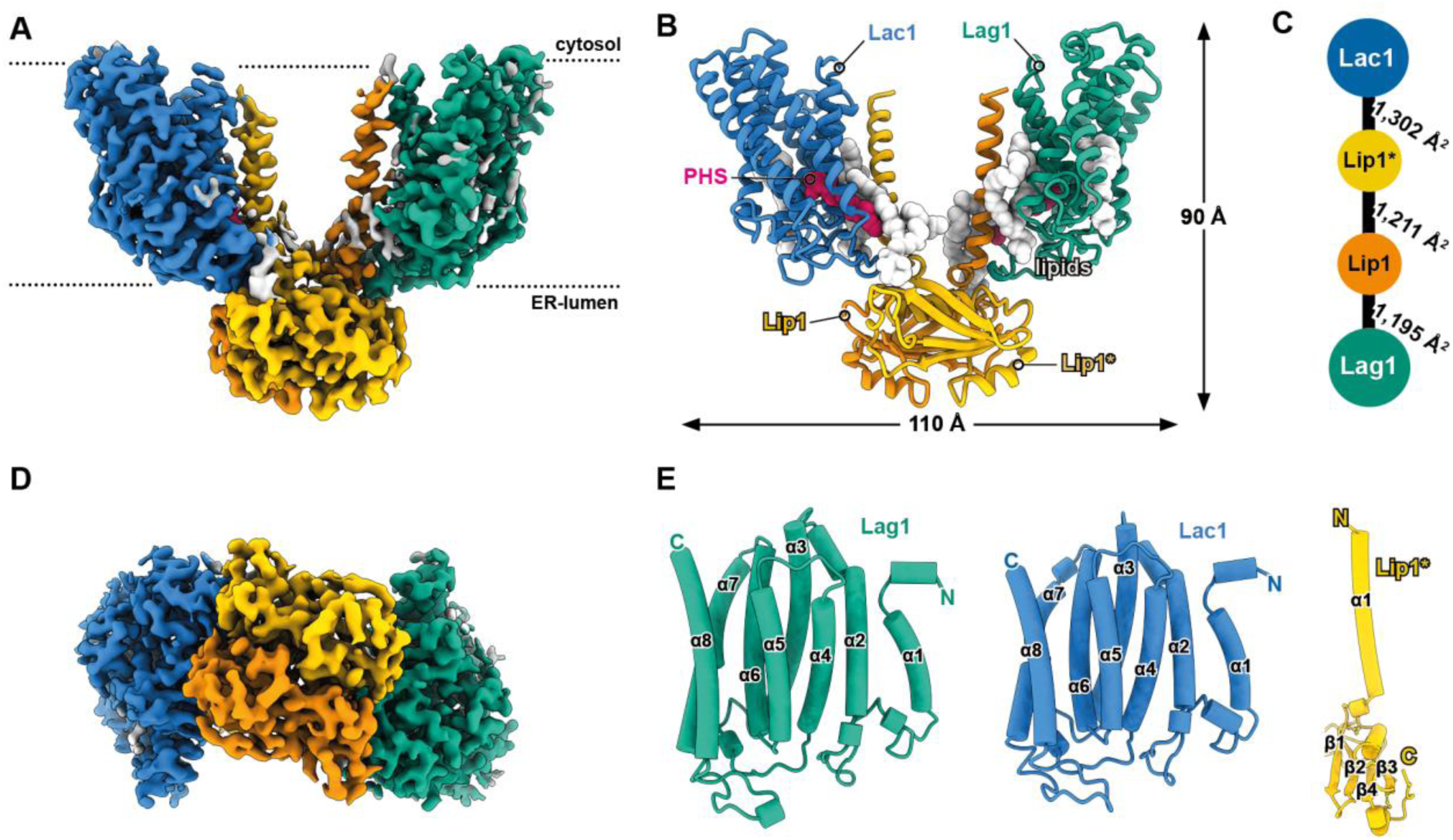
Architecture of the yeast ceramide synthase complex. **A** Cryo-EM reconstruction of the yeast CerS complex at 3.0 Å represented from the side relative to the membrane plane. **D** A view from the ER lumen is given. **B** Atomic model shown as a cartoon representation of the complex with the enzymatic core-subunits Lac1 (blue) and Lag1 (green). Lip1 (orange) and Lip1* (yellow) are the small-regulatory subunits. Phytosphingosine molecules (PHS, pink) and several lipids (white) are shown as spheres. **C** Interaction diagram of the CerS subunits and their respective contact areas in Å^2^. **E** Topology representation of the CerS subunits with eight transmembrane helices (α1-8) for Lag1/Lac1 and one helix in Lip1.

The presence of two disulfide bonds between S142-S102 and S75-S53 in each Lip1 subunit reveals the overall orientation of the complex, with the Lip1-Lip1 positioned in the ER lumen (Fig. 2B). Consequently, the N-terminus of Lip1 resides in the cytosol. The first 18 amino acids of Lip1 are not resolved in the structure, potentially due to the flexibility of this region (Sup. Fig. 8B). Each Lip1 interacts with one of the respective catalytic subunits Lag1/Lac1 through their transmembrane helices. Attachment is driven by the hydrophobic interactions between the transmembrane helices TM1 from Lip1 and TM5-6 from Lag1/Lac1 (Fig. 2C-F). In addition, a small interface exists between loop regions in Lip1 (aa 106-124) and Lag1 (aa 332 −342). Notably, the positioning of the two catalytic subunits disallows a direct interaction of the catalytic subunits via the proposed dimerization motif identified in mammalian cells^36^ (Sup. Fig. 7). Despite their high sequence identity of 68.6 %, our structural data enabled unambiguous assignment and differentiation of Lag1 and Lac1 in our density maps (Sup. Fig. 6D, E), confirming that Lac1 and Lag1 form a heterodimer within the CerS complex. Since Lac1 is also co-purified by performing pulldowns of GFP-tagged Lag1, this further supports the essential role of Lip1 in the complex (Sup. Fig. 1J). While the DxRSDxE motif (aa 380-386 in both Lag1 and Lac1) are not resolved in our structure, the C-termini of Lag1 and Lac1 are widely spaced and are unlikely to interact without major conformational or environmental changes (Sup. Fig. 7).

**Fig. 2:**
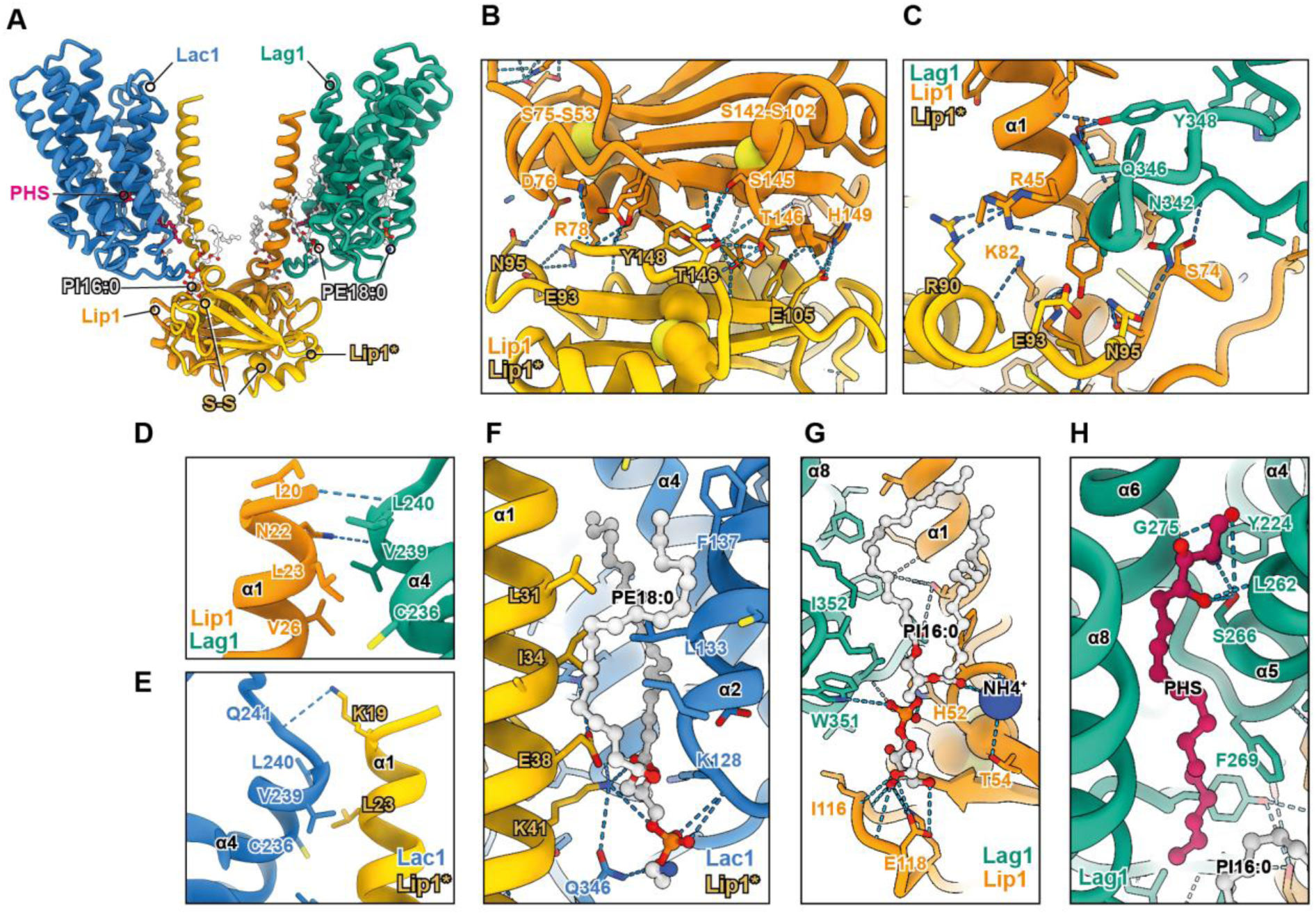
Subunit and small-molecule interactions within the ceramide synthase. **A** Atomic model in cartoon-representation of the different subunits of the CerS with Lac1 in blue, Lag1 in green, Lip1 in orange, and Lip1* in yellow. Phytosphingosine (PHS) in pink and the lipids PE18:0 and PI16:0 in white are shown in ball-and-stick representation. **B** Interaction-interface of Lip1-Lip1*. Disulfide bonds between S142-S102 and S75-S53 are shown as spheres. **C** Binding of the three subunits Lag1, Lip1, and Lip1* at the luminal side of the complex. **D** Binding site of Lip1 and Lag1 on the cytosolic side of the complex. **E** Binding of Lip1* and Lac1 near the cytosol. **F** PE18:0 bridges the binding of Lac1 and Lip1* in the luminal membrane leaflet. **G** PI16:0 and ammonium bridge the binding of Lip1 and Lag1 within the luminal membrane leaflet. **H** PHS-binding site with residues from Lag1. All subunits are color-coded consistently. Dashed lines represent polar contacts, and residues without direct lines illustrate hydrophobic interactions.

The overall fold of the catalytic subunits resembles the highly conserved TRAM/LAG1/CLN8 homology domain harboring the proposed six transmembrane helix architecture (TM3-TM8; Fig. 1E, Sup. Fig. 6A, B). Two additional transmembrane helices (TM1-TM2) at the N-terminus of each catalytic subunit place both the N- mand the C-terminus of Lag1/Lac1 into the cytosol (Fig. 1E, Sup. Fig. 6A, B). This orientation of the complex is in line with the previously reported phosphorylation sites (S23 and S24 by Ypk1 and S393, S395, and S397 by CK2^34,35^). Both termini of the catalytic subunits Lag1 and Lac1 (amino acids 1-74 and 384-411) are not resolved in the structure (Sup. Fig. 6D, Sup. Fig. 7A).

### Several lipids are co-purified with the ceramide synthase complex

Besides the well-resolved trans-membrane core of the CerS, we also identified several lipids co- purified with the complex. Each catalytic subunit shows two densities, best fitted by two glycerol- phospholipid molecules with small headgroups, most likely phosphatidylethanolamine (Fig. 2F, Sup. Fig. 4H,5C). Both headgroups point towards the luminal site of the ER bilayer. Another lipid density is located at the luminal site of the complex between the Lip1 luminal domain and a loop of the catalytic subunits. The prominent headgroup suggests that this density represents a molecule of phosphatidylinositol (PI, Fig. 2G, Sup. Fig. 4G, 5G). Peculiarly, the PI molecule sticks out of the ER lumen-facing membrane plane (Fig. 3A, B). While the inositol headgroup is near E118 in a loop of Lip1, the phosphate moiety of the PI is located proximal to H52 of Lip1. Interestingly, the glycerol backbone and the first carbon atoms of the acyl chains are surrounded by hydrophobic amino acids in luminal loop regions from Lip1 (W50, F51) and Lag1 (L341, F343). Together, this suggests that the PI contributes to stabilizing the interaction of Lip1 with either catalytic subunit.

**Fig. 3:**
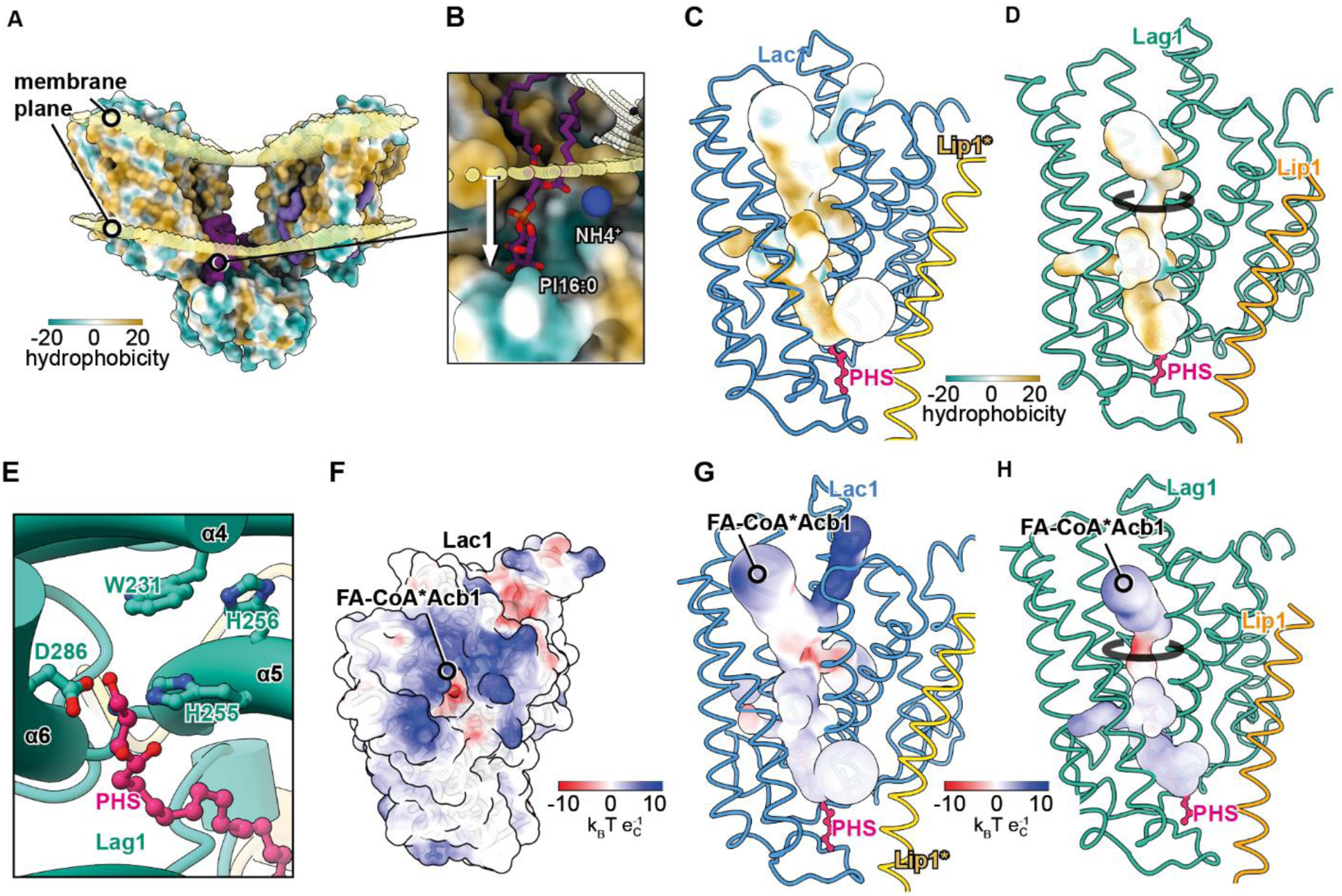
Surface properties of the ceramide synthase linked to its function. **A** Surface representation of the CerS complex color-coded by its hydrophobicity. The curved membrane plane (transparent yellow) was calculated by PPM3 using the yeast ER-membrane model. PI16:0 is shown as a dark violet surface. **B** Focused view from (A) onto PI16:0. Displacement of PI16:0 (stick representation) from the luminal membrane leaflet. **C** Internal cavities within Lac1 color-coded by hydrophobicity, partially occupied by a PHS molecule (pink). **D** Internal cavities within Lag1 color-coded by hydrophobicity, partially occupied by a PHS molecule (pink). **E** Catalytic site of the CerS with the catalytic residues W231, H256, H255, and D286 in Lag1. The PHS molecule (pink) is positioned close to these residues. **F** Surface- representation of Lac1 color-coded by its electrostatic potential. A positive patch on the cytosolic side is annotated as putative fatty acid-coenzyme A (FA-CoA) and Acb1 binding region. **G, H** Internal cavities within Lac1 (G) and Lag1 (H) are color-coded by the electrostatic potential, which is partially occupied by a PHS molecule (pink). The putative FA-CoA*Acb1 entry site is annotated.

Another density is located in a hydrophobic cavity at the center of the complex in the luminal part of the lipid bilayer. It resembles the structure of a lyso-lipid with only one acyl chain and a very small headgroup (Fig. 2H, Sup. Fig. 4F, 5C). Since CerSs use a sphingoid base as a substrate, this likely reflects a molecule of PHS. Indeed, the presence of a molecule with a mass of 318.4999 m/z and its 60 m/z transition mass in the purified complex could be confirmed by mass spectrometry (Sup. Fig. 1H).

### Ceramide synthase harbors a large cavity carrying the active site

The catalytic subunits Lag1 and Lac1 harbor two specific TM bundles (TM3–5, TM6–8) tilting away from each other, generating space in the center for the catalytic chamber (Fig. 4G, Sup. Fig. 6C). TMs 7 and 8 are two long, diagonal TM segments with 30 amino acids each. TM2 tilts similarly to the TM 6-8 helix bundle and completes the chamber. This cavity is accessible from the cytoplasmic site and the intermembrane space towards the direction of the Lip1 scaffold (Fig. 3C, D).

**Fig. 4:**
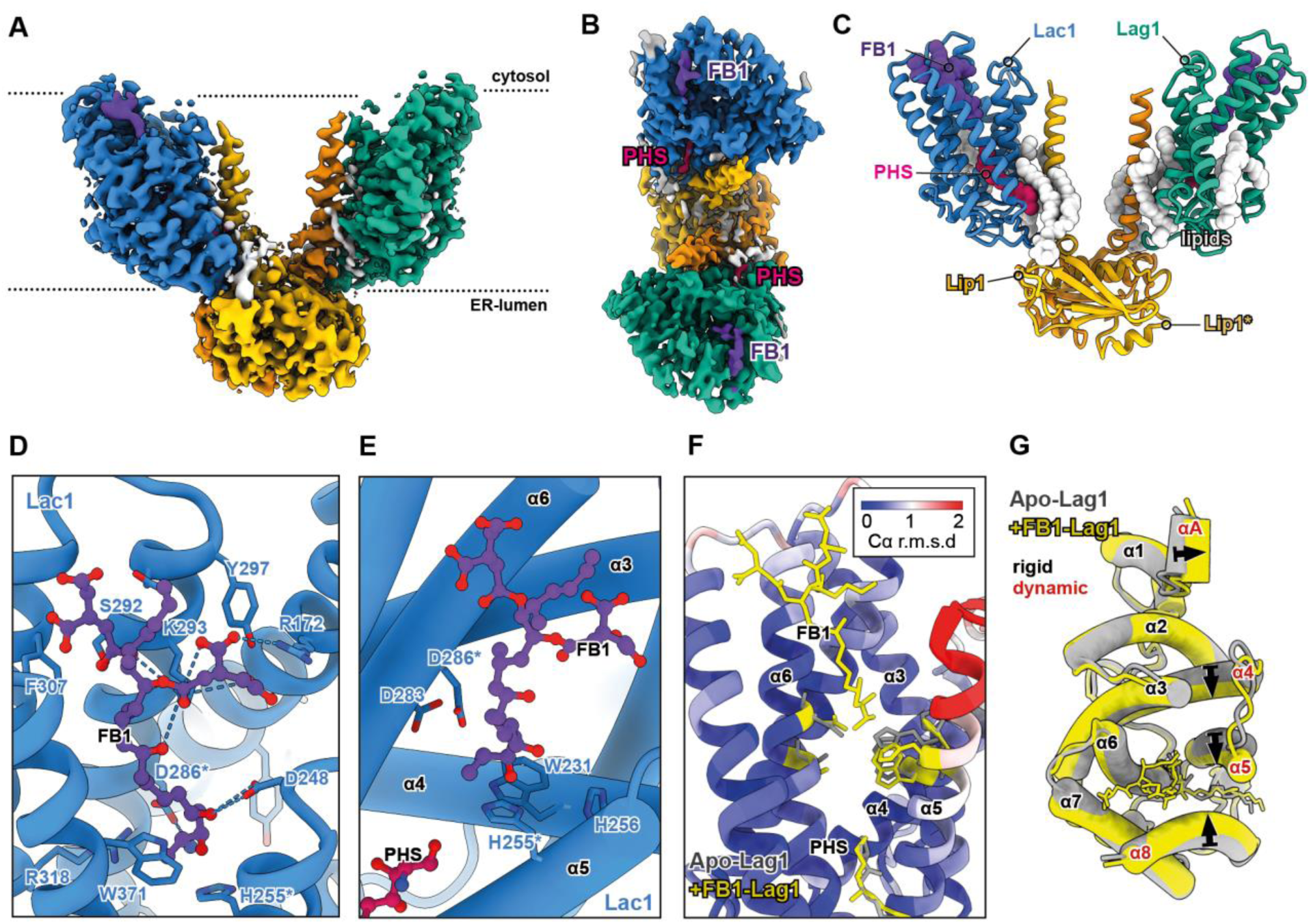
FB1 blocks the cytosolic ligand entry site of the ceramide synthase. **A** Cryo-EM reconstruction of the fumonisin B1 (FB1) bound CerS at 3.2Å. A part of Lac1 TM8 was omitted to show the FB1 density (violet). **B** Rotated view of the cryo-EM density viewing from the cytosol to reveal the FB1-binding to Lac1 and Lag1. PHS (pink) is present in the FB1-bound CerS. **C** Cartoon-representation of the CerS subunits with Lac1 in blue, Lag1 in green, Lip1 in orange, and Lip1* in yellow. Sphere- representation of FB1 (violet), PHS (pink), and lipid molecules (gray). **D** Binding site of FB1 (ball-and-stick representation) with Lac1. Polar contacts are highlighted with blue dashed lines. Residues with hydrophobic interactions are shown without lines. Residues H255 and D286 are annotated with an asterisk to highlight the overlapping interaction with FB1 and their catalytic function. **E** FB1 and PHS are positioned near the catalytic residues of Lac1 within TM4-6. **F** Structural comparison of CerS apo (gray) and the FB1-bound structure (yellow). The cartoon representation is color-coded by the Cα (root-mean-square deviation, r.m.s.d.). FB1, PHS, and the catalytic residues are shown as sticks. **G** Structural comparison of the CerS apo (gray) and FB1-bound Lag1 (yellow). FB1-binding induces a shift within the Lip1-facing transmembrane-helices 4,5 and 8, and a small helix denoted αA. The membrane-exposed TM1,2,3,6,7 largely remain rigid.

Computational studies on the AlphaFold predicted structure of the human CerS 5 (CerS5) suggested a funnel-like crevice lined with positively charged amino acids at the cytosolic site^38^. The corresponding residues (K171 and R172 in Lag1 and Lac1) are also found at the cavity’s entrance facing the cytosol. Similarly, previous studies have identified several H, D, and W residues critical for the catalytic activity of the CerS (Fig. 3F). The evolutionarily conserved residues (H255, H256, D283, D286, and W371 in Lag1 and Lac1), which are critical for catalytic activity, are located at the midpoint of the cavity, supporting the location of the active center of the enzyme (Fig. 3E). Interestingly, the sphingoid base, a substrate for the CerS, is located further down in the funnel-like cavity, potentially reflecting a pre-loaded state of the CerS (Fig. 2H). The free amino group of the base that is N-acylated by the enzyme would have to move up in the funnel upon binding the fatty acid CoA, requiring conformational changes in the enzyme.

### Fumonisin B1 blocks the active center of the ceramide synthase from the cytoplasmic site

To test if the enzyme can undergo conformational changes and explore the mechanism of inhibition, we determined the fumonisin B1 bound structure of the CerS. Fumonisin B1 is a known inhibitor of the CerS. It was proposed that the structural similarities between fumonisins (1-deoxy- analogs of sphinganine; Sup. Fig. 1K) and sphingoid bases enable inhibition of CerS^40^. Single particle cryo-EM of the FB1-treated CerS resulted in a structure resolved with 3.2 Å resolution (Fig. 4, Sup. Fig. 3, Sup. Tab. 1). Compared to the uninhibited structure, large-scale conformational changes were not observed. However, an additional density appeared at the cytosolic entrance of the funnel-like cavity (Fig. 4A, B). The data show that the sphinganine-like hydrocarbon chain of the fumonisin B1 extends into the hydrophobic cavity, placing the amino group near the H, D, and W residues important for the catalytic activity (Fig. 4D, E; Sup. Fig. 5E, F; Sup. Fig. 8A; Sup. Fig. 9). The ester-linked tricarballylic acids of FB1 are blocking the entrance of the cavity. Together, this explains the inhibitory function of FB1 on CerSs.

Interestingly, the binding of FB1 also resulted in an opening of a hydrophobic cavity facilitated through motions in TM4 and 5 (Fig. 4F, G). Despite the inhibition, the same long-chain base substrate is present in the cavity (Fig. 4B, C). Its overall orientation and position did not change. Since binding fumonisin B1 to the cytosolic entrance led to conformational changes, binding the specific acyl-CoA substrate will likely also cause a conformational change. This could then result in the correct positioning of the substrates in the active center of the enzyme.

## Discussion

Ceramides are crucial metabolic intermediates in sphingolipid metabolism and can act as signaling molecules. In addition, ceramides are fundamental to many biological processes, including cancer development, obesity, type 2 diabetes mellitus, skin barrier function, and lipid storage. Dysregulation can lead to the metabolic syndrome or neurodegenerative disorders. Understanding how CerSs catalyze the ceramide biosynthesis reaction and how fatty acids are selected is crucial to developing selective therapeutics that can potentially circumvent the catastrophic effects of global ceramide inhibition. Structural data that could explain the molecular mechanism of these essential enzymes is absent. Here, we report the first structures of the yeast CerS as a sphingoid base-bound structure and a fumonisin B1 inhibited complex.

The yeast CerS is a heterotetrametric complex exhibiting a pseudo-C2 symmetry. At its central axis, a homodimer of Lip1 acts as a scaffold to which the two catalytic subunits Lac1 and Lag1 are bound. While we did not observe homo-tetramers containing either two copies of Lag1 or Lac1, respectively, the architecture of the complex would, in principle, allow their presence. This could explain why only the double deletion of *LAG1* and *LAC1* causes severe effects^32^. In contrast, the Lip1 subunit is essential for yeast cells^33^. Since Lip1 does not participate in forming the active site, we propose that it functions as a scaffold protein required for the activity of the CerS. Lip1 could establish a specific membrane environment required for substrate loading of the CerS. Along this line, we find an opening of the Lag1/Lac1 cavity in the intermembrane space towards the Lip1 transmembrane helices. In our structure, the cavity is occupied by a density that fits one of the CerS substrates, a sphingoid base. A computational study on a predicted structure of the mammalian CerS5 also predicted an opening of the cavity in the intermembrane space, and the authors speculate that this is important for substrate loading and the exit of ceramide^41^. So far, no Lip1 homolog has been identified in mammalian cells.

In general, structures of other lipid synthesizing enzymes resemble some features we also detect in the structure of the CerS. These include large (>28 aa) transmembrane helices tilted in the membrane and forming the active site. Similar architectures have been observed in the structures of the diacylglycerol acyl transferases (DGATs) and membrane-bound O-acyltransferases (MBOATs)^42,43^. In addition, these enzymes also have intra-bilayer opening cavities required for loading the hydrophobic lipid substrates into the active site of the enzymes.

Lag1 and Lac1 both harbor eight transmembrane helices placing both termini in the cytosol^44^. Therefore, the yeast CerS appears to differ from the mammalian CerSs, which harbor seven trans-membrane helices. Additionally, yeast CerSs lack the cytosolic homeobox found in the catalytic subunits of mammalian CerSs 2-6^31^. Furthermore, in mammals, the elongated N-terminus that forms the first two transmembrane domains in yeast is absent. The N-terminus of Lip1 and both termini of the catalytic subunits face the cytosol, which aligns with previously identified phosphorylation sites. Lip1 is phosphorylated at residues S2 and T5^45^, while Lag1 and Lac1 are phosphorylated at both termini by the yeast Ypk kinases or the casein kinase 2 (CK2)^46,47^. One of the phosphorylation sites is found in the DxRSDxE motif in the C-terminus of Lag1/Lac1. The sequence is highly conserved and acts as a dimerization region for mammalian CerSs^48^. Although not resolved in our structure, presumably due to flexibility, the overall architecture of the complex indicates that both DxRSDxE motifs point in different directions, which prohibits any physical interactions. Therefore, the DxRSDxE motif could be responsible for forming higher-order interactions between several CerS complexes in yeast. Potential higher-order complexes in yeast cells for Lag1 and Lac1 have already been observed in hydroxyurea- treated cells^49^.

Our fumonisin B1-containing structure resolves the mode of action for the CerS inhibitor. We resolved bound fumonisin B1 in the cytosolic site of the large cavity of Lag1/Lac1. It has been suggested that fumonisin B1 resembles both parts of the fatty acid CoA molecule and the sphingoid base^50^. The amino group of FB1 at the second carbon atom is close to the putative active site of the CerS containing H255/256, D283/286, and W231 (Sup. Fig. 6d). The two-ester linked tricarballylic acids have been proposed to mimic the two phosphate groups of the CoA moiety of an acyl CoA. However, the fumonisin B1 does not interact with the positively charged lysins and arginines proposed for CoA binding in the computational studies of human CerS5^41^. Nevertheless, the binding of fumonisin B1 appears sufficient to occupy the binding pocket and prevent the acyl-CoA from binding and thus inhibit enzyme activity.

In summary, we present the first structure of a CerS and provide a testable explanation for the substrate loading of the enzyme. In the future, these structures will help to identify the molecular mechanism of substrate loading and the catalytic mechanism of the enzymes. The high conservation of the catalytic domain can also be a starting point to develop subunit-specific inhibitors for the different CerSs analogous to the inhibitors for CerS1^51^. Selective therapeutics will be necessary to circumvent the potentially catastrophic effects of global ceramide inhibition.

## Material and Methods

### Yeast strains

Yeast strains and plasmids used can be found in Supplementary Tables 2 and 3. For purifications, the inducible GAL1 promoter was genomically integrated upstream of LAG1, LAC1, and LIP1, including a 3xFLAG tag inserted at the N-terminus of LIP1^52^.

### Purification of 3xFLAG tagged CerS from *S. cerevisiae*

Purification of 3xFLAG tagged ceramide synthase was performed as previously reported for the yeast SPOTS complex^53^. For this, yeast cells were grown for 24 h in a yeast peptone (YP) medium containing 2% galactose (v/v). Cells were harvested, washed in lysis buffer (50mM HEPES-KOH (pH 6.8), 150mM KOAc, 2mM MgOAc, 1mM CaCl2, 200mM sorbitol), and resuspended in lysis buffer supplemented with 1mM phenylmethylsulfonylfluoride (PMSF) and 1x FY protease inhibitor mix (Serva) in a 1:1 ratio (w/v). Resuspended cells were frozen dropwise in liquid nitrogen and pulverized in 15 cycles of 2 mins at 12 CPS using a 6875D Freezer/Mill Dual-Chamber Cryogenic Grinder (SPEX SamplePrep). After the yeast powder was thawed in lysis buffer containing 1mM PMSF and 1x FY, cell debris was removed at 1,000 g at 4 °C for 20 mins. Microsomal membranes were collected at 44,000 g at 4 °C for 30 mins, resuspended in lysis buffer, and then diluted with IP buffer (50mM HEPES-KOH (pH 6.8), 150mM KOAc, 2mM MgOAc, 1mM CaCl2, 15% glycerol) supplemented with 1% glyco-diosgenin (GDN) and protease inhibitors. After solubilization for 1.5 hours at 4 °C and a centrifugation step at 44,000 g at 4 °C for 30 mins, the supernatant was added to α-FLAG resin (Sigma Aldrich). Beads were washed twice with IP buffer containing 0.1 % and 0.01 % GDN, respectively, after incubation for 45 mins at 4 °C. 3xFLAG peptide was added twice and incubated together with α-FLAG beads for 45 mins and 5 mins on a turning wheel. Proteins were eluted at 460 g at 4 °C for 30 s and concentrated in a 1% GDN pre-equilibrated 30 kDa Amicon Ultra centrifugal filter (Merck Millipore). The concentrated eluate was spun through a Zeba Spin Desalting Column 40 kDa MWCO (Thermo Fisher Scientific) to remove high glycerol concentrations and then directly used for Cryo-EM.

### Proteomics

Proteomics analysis was performed as described previously^54^. Purified proteins were treated according to the iST Sample Preparation Kit (PreOmics) protocol. Dried peptides were resuspended in LC-Load and used to perform reversed-phase chromatography on a Thermo Ultimate 3000 RSLCnano system connected to a QExactivePLUS mass spectrometer (Thermo Fisher Scientific) through a nano-electrospray ion source. Peptides were separated on a 50-cm PepMap C18 easy spray column (Thermo Fisher Scientific) with an inner diameter of 75 µm and a column temperature of 40°C. The peptides were eluted from the column via a linear gradient of acetonitrile from 12 to 35% in 0.1% formic acid for 80 min at a constant flow rate of 250 nl/min followed by a 20 min increase to 60%, and finally, 10 min to reach 90% buffer B. Eluted peptides from the column were directly electro sprayed into the mass spectrometer. Mass spectra were acquired on the QexactivePlus in a data-dependent mode to automatically switch between full scan MS and up to ten data-dependent MS/MS scans. The maximum injection time for full scans was 50 ms, with a target value of 3,000,000 at a resolution of 70,000 at m/z 200. The 10 most intense multiply charged ions (z ≥ 2) from the survey scan were selected with an isolation width of 1.6 Th and fragments with higher energy collision dissociation (Olsen et al., 2007) with normalized collision energies of 27. Target values for MS/MS were set at 100,000 with a maximum injection time of 80 ms at a resolution of 17,500 at m/z 200. The dynamic exclusion of sequenced peptides was set at 20 s to avoid repetitive sequencing. The resulting MS and MS/MS spectra were analyzed using MaxQuant (version 2.4.4.0, www.maxquant.org/)^55,56^. The resulting data were analyzed using Perseus (V2.0.11, www.maxquant.org/perseus)^57^. Significance lines in the volcano plot of the Perseus software package corresponding to a given FDR were determined by a permutation-based method^58^. The mass spectrometry proteomics data have been deposited to the ProteomeXchange Consortium via the PRIDE partner repository^59^.

### Size-exclusion chromatography

The concentrated eluate of the ceramide synthase was applied to a Superose 6 Increase 5/150 column (Cytiva) with 50 µl elution volume using ÄKTA go purification system (Cytiva). IP buffer with 0.008% GDN and no glycerol was used for elution. Peak fractions were collected, and the presence of all three subunits, Lag1/Lac1/Lip1, was confirmed by mass spectrometric analysis.

### GFP-Trap pulldown experiments

For pulldown experiments, cells were grown in 50 ml YPD medium inoculated from an overnight pre-culture and grown to exponential growth phase at 30 °C. The same amount of cells were harvested at 2,272 g at 4°C for 5 min, washed with ice-cold water, and snap-frozen as pellets in liquid nitrogen. Cells were lysed with 500 μl acid-washed glass beads in 500 μl GFP-Trap pulldown buffer (20 mM HEPES, pH 7.4, 150 mM KOAc, 5% glycerol, 1% GDN, Roche Complete Protease Inhibitor Cocktail EDTA free, Roche) using a FastPrep (MP biomedicals). Cell lysates were spun down at 1,000 *g* at 4°C for 10 min, followed by an incubation at 4 °C rotating for 30 min. Supernatants were centrifuged at 21,000 g at 4 °C for 10 min and then incubated for 10 min at 4 °C with 12.5 μl pre-equilibrated GFP-Trap beads (Chromotek) rotating. Beads were washed twice with GFP-Trap pulldown buffer and then four times with washing buffer (20 mM HEPES pH 7.4, 150 mM KOAc, 5% glycerol). Bound proteins were digested and treated according to the iST Sample Preparation Kit (PreOmics) protocol. Dried peptides were resuspended in LC-Load, and reversed-phase chromatography was performed on a Thermo UltiMate 3000 RSLCnano system connected to a TimsTOF HT mass spectrometer (Bruker Corporation, Bremen) through a Captive Spray Ion source. Separation of peptides was conducted on an Aurora Gen3 C18 column (25 cm x 75um x 1.6um) with a CSI emitter (Ionoptics, Australia) at 40 °C. Peptides were eluted from the column via a linear gradient of acetonitrile from 10–35 % in 0.1 % formic acid for 44 min at a constant flow rate of 300 nl/min following a 7 min increase to 50 %. Finally, 4 min to reach 85 % buffer B. Eluted peptides were directly electro-sprayed into the mass spectrometer at an electrospray voltage of 1.5 kV and 3 l/min Dry Gas. Positive ion polarity with a MS range from 100 to 1700 m/z was used as the MS settings of the TimsTOF. The resulting data were analyzed and visualized using MaxQuant (V2.4.9.0, www.maxquant.org)^55,56^ and Perseus (V2.0.11, www.maxquant.org/perseus)^57^. Significance lines in the volcano plot of the Perseus software package corresponding to a given FDR were determined by a permutation-based method. The mass spectrometry proteomics data have been deposited to the ProteomeXchange Consortium via the PRIDE partner repository^59^.

### Expression and purification of yeast Acyl-CoA binding protein 1 (Acb1)

An N-terminally glutathione S-transferase (GST)-tagged version of yeast Acyl-CoA binding protein 1 (Acb1) was transformed in *Escherichia coli* BL21 Rosetta cells using a pGEX-6P-1 expression vector system. Cells were grown in Lysogeny broth (LB) media to an OD600 of 0.6, cooled down on ice for 1h, and protein expression was induced by adding 0.25 mM IPTG overnight at 16°C and 250rpm. Cells were harvested, washed, and resuspended in a lysis buffer (50mM Tris-HCl pH 7.4, 300mM NaCl). Cells were lysed in a Microfluidizer (Microfluidics, Westwood, MA, USA), and the lysate was clarified by centrifugation at 17,000 g for 20 min at 4°C. After an incubation for 2h on a nutator with 1.5 ml pre-equilibrated Glutathione Sepharose 4B resin (Cytiva, 17075601), beads were washed thoroughly, and Acb1 was eluted in elution buffer (25mM Tris pH 7.4, 120mM NaCl, 1mM DTT) containing 20mM reduced glutathione. Proteins were dialyzed twice against 2L elution buffer using a dialyzing tube with 3.5 kDa molecular weight cut-offMWCO (Spectra/Por). Protein purity was confirmed by SDS-PAGE and mass spectrometric analysis as described before.

### LCBs analysis of purified ceramide synthase

LCBs were extracted from 20 µg of purified protein in three technical triplicates and measured (n=3) as described previously^53^. Briefly, 150 mM ammonium formate was added to purified proteins, and as an internal standard, sphingosine d17:1, Avanti was added. Lipids were extracted with 2:1 chloroform/methanol as described previously^60,61^. Dried lipid films were dissolved in a 65:35 (v/v) mixture of Buffer A (50:50 water/acetonitrile, 10 mM ammonium formate, and 0.1% formic acid) and B (88:10:2 2-propanol/acetonitrile/water, 2 mM ammonium formate and 0.02% formic acid). Phytosphingosine 18:0 (PHS; Avanti Polar Lipids) was used to prepare an external standard curve. Samples were analyzed on an Accucore C30 LC column (150 mm × 2.1 mm 2.6 µm Solid Core; Thermo Fisher Scientific) connected to a Shimadzu Nexera HPLC system and a QTRAP 5500 LC-MS/MS (SCIEX) mass spectrometer. For the gradient, 40% B was used for 0.1 min, which was then increased to 50% over 1.4 min, followed by an increase to 100% over 1.5 min. 100% B was kept for 1 min and decreased to 40% B for 0.1 min. 40% B was maintained until the end of the gradient. A constant flow rate of 0.4 ml/min was used with a total analysis time of 6 min and an injection volume of 1 or 2 μl. The MS data were measured in positive ion, scheduled MRM mode without detection windows (Sup. Tab. 6).

### Cryo-EM sample preparation and data collection

Prior to cryo-EM, the sample quality was analyzed by negative-stain electron microscopy, as previously described^62^ (Sup. Fig. 1G). Micrographs were manually collected on a JEM2100plus transmission electron microscope (Jeol) operating at 200 kV and equipped with a Xarosa CMOS (Emsis) camera at a nominal magnification of 30,000, corresponding to a pixel size of 3.12 Å per pixel.

For cryo-EM ligand-binding studies, 8.5 mg/ml CerS was incubated for one hour on ice with 100 µM FB1. Excess glycerol was removed directly before freezing using a 40 kDa MWCO Zeba Micro spin-desalting column (Thermo Fisher). C-flat grids (Protochips; CF-1.2/1.3-3Cu-50) were glow-discharged using a PELCO easiGlow device at two consecutive runs of 15 mA for 45 s. A sample volume of 3 µL was applied to a grid immediately before plunge freezing. The samples were vitrified in a Vitrobot Mark IV (Thermo Fisher) at 100% relative humidity at 4 °C. All datasets were collected on a Glacios microscope (Thermo Fisher) operating at 200 kV and equipped with a Selectris energy filter (Thermo Fisher) set to a slit width of 10 eV. Movies were recorded using a Falcon 4 direct electron detector (Thermo Fisher) at a nominal magnification of 165,000, corresponding to a pixel size of 0.68 Å per pixel. Movies were saved in the electron-event representation (EER) format and recorded at a total dose of 50 e− per Å^2^. All datasets were collected automatically using the EPU software (v.3.3, Thermo Fisher) with a defocus range of −0.8 to −2.0 μm.

### Cryo-EM data processing

All datasets were processed using cryoSPARC (v.4.2). The corresponding processing workflows are shown in supplementary Fig. 2-3. Movies were preprocessed with patch-based motion correction, patch-based CTF estimation and filtered by the CTF fit estimates using a cutoff at 4.5 or 5.0 Å in cryoSPARC live.

Template-based particle picks from live data processing were used for 2D classification. Class averages with recognizable secondary structures were used for training TOPAZ. The resulting particle picks from TOPAZ were classified in 2D, and well-aligning particles were subjected to multi-class ab initio reconstruction. Subsequently, good reconstructions from a heterogeneous refinement of the four-fold Fourier-cropped particle images were combined and subjected to non-uniform refinement. A wide mask, including the detergent micelle, was used, and auto-mask tightening was disabled. Re-centered particles were re-extracted at the full box of 450 px. Two rounds of heterogeneous refinement and non-uniform refinement were used to reduce heterogeneity. In the case of the apo dataset, the CTF-fit cutoff was set to 4.5 Å. 3D classification without alignment was used in PCA-mode initialization, two starting classes, a filter resolution of 3 Å, and a tight mask around the CerS. Finally, local refinement with a provided tight mask resulted in an overall resolution of 3.0 Å for the CerS in the apo state and 3.2 Å for the FB1 / C26CoA-incubated reconstruction.

Details about data collection for individual datasets and validation statistics are given in sup. Tab. 1. Reported B-factors resulted from unsupervised auto-sharpening during refinement in cryoSPARC. To aid model building, unsharpened half-maps, FSC-based resolution, and the molecular mass were supplied as input for density modification within Phenix *phenix.resolve_cryo_em*^63^.

### Model building, refinement, and validation

The initial atomic model of the CerS complex was predicted with AlphaFold 2 multimer^64^ and fitted into the densities as rigid bodies in ChimeraX^65^. The structure was manually inspected using Coot (v.0.9)^66^ and ISOLDE^67^. Models were refined using *phenix.real_space_refine* within Phenix (v.1.19)^68^. Validation reports were automatically generated using MolProbity^69^ within Phenix.

### Fluorescence-based enzymatic assay

To measure CerS activity, CoA-SH released during the catalyzed reaction was monitored using the Coenzyme A Assay Kit (MAK034, Sigma-Aldrich). All assays were performed on a 200 μl scale in an IP buffer with 0.01% GDN, 40 µM phytosphingosine (PHS), and 250 µM C26-CoA. If indicated, 0.3 % BSA or 0.74 µg Acb1 were included in the reaction buffer. As a control, Fumonisin B_1_ (FB_1_), a well-established inhibitor of CerS activity, was added to a final concentration of 30 µM. To all other samples, Dimethyl sulfoxide (DMSO) was added since FB1 was dissolved in DMSO. The reactions were initiated by adding 1 µg purified protein and incubated at RT for 1 h. Afterward, the samples were deproteinized using a 10 kDa MWCO concentrator (Merck Millipore) and treated in a 96-well plate according to the Coenzyme A Assay Kit protocol (Sigma-Aldrich). The fluorescence of the fluorescent product was measured in a SpectraMax iD3 Multi-Mode microplate reader (λ_Ex_ = 535 nm/λ_Em_ = 587 nm). Measured fluorescence intensities of protein-free samples were subtracted from the fluorescence intensities of the protein-containing samples.

## Data availability

All density maps and models were deposited in the Electron Microscopy Data Bank and the Protein Data Bank. The PDB IDs are 8QTN for the *Sc*CerS apo and 8QTR for the FB1-bound *Sc*CerS. The respective EMDB IDs are EMD-18652 and EMD-18653. All structural data was visualized with ChimeraX, and protein interactions were analyzed with the help of PDBe PISA^70^. 2D ligand-protein interaction diagrams were calculated using LigPlot+^71^.

## Analysis of biomacromolecular channels

Tunnels and interior cavities were mapped using MOLE 2.5. The following parameters were used: bottleneck radius:1.2 Å, bottleneck tolerance, 3 Å; origin radius, 5 Å; surface radius, 10 Å; probe radius, 5 Å; interior threshold 1.1 Å^72^. The physicochemical properties of the MOLE output were visualized in ChimeraX.

## Acknowledgments

We thank members of the Fröhlich and Moeller labs for their valuable comments on the manuscript. This work was funded by the DFG (SFB1557 P6 to FF, P11 to AM, FR 3647/2-2 and FR 3647/4-1 to FF and INST.190/196-1 FUGG, MO2752/3-6 and SPRUNG Stay inspired 15- 76251-2-04/22 (10743/2022) to AM), BMBF 532 01ED2010 to AM. JHS is supported by a fellowship from the Friedrich Ebert foundation.

## Author contributions

Investigation: JHS, LC, CK, VNW, BME, SW. Formal analysis: JHS, LC, CK, VNW, BME, SW, isualization: JHS, LC, CK, FF. Conceptualization: FF, AM, DJ. Funding acquisition: FF, AM. Writing original draft: JHS, LC, CK, FF, AM. Writing review and editing: JHS, LC, CK, FF, AM.

