## Supplementary-Information for "Structure of the yeast ceramide synthase"

[1] Osnabrück University
Department of Biology/Chemistry
Structural Biology section
49076 Osnabrück, Germany

[2] Osnabrück University
Department of Biology/Chemistry
Bioanalytical Chemistry section
49076 Osnabrück, Germany

[3] Osnabrück University
Center of Cellular Nanoanalytic Osnabrück (CellNanOs)
49076 Osnabrück, Germany

[†] These authors contributed equally to this work

[#] For correspondence:
 (F. F.); (A. M.);

**Keywords:** ceramide synthase, ceramide, sphingolipids, very-long chain fatty acid CoA

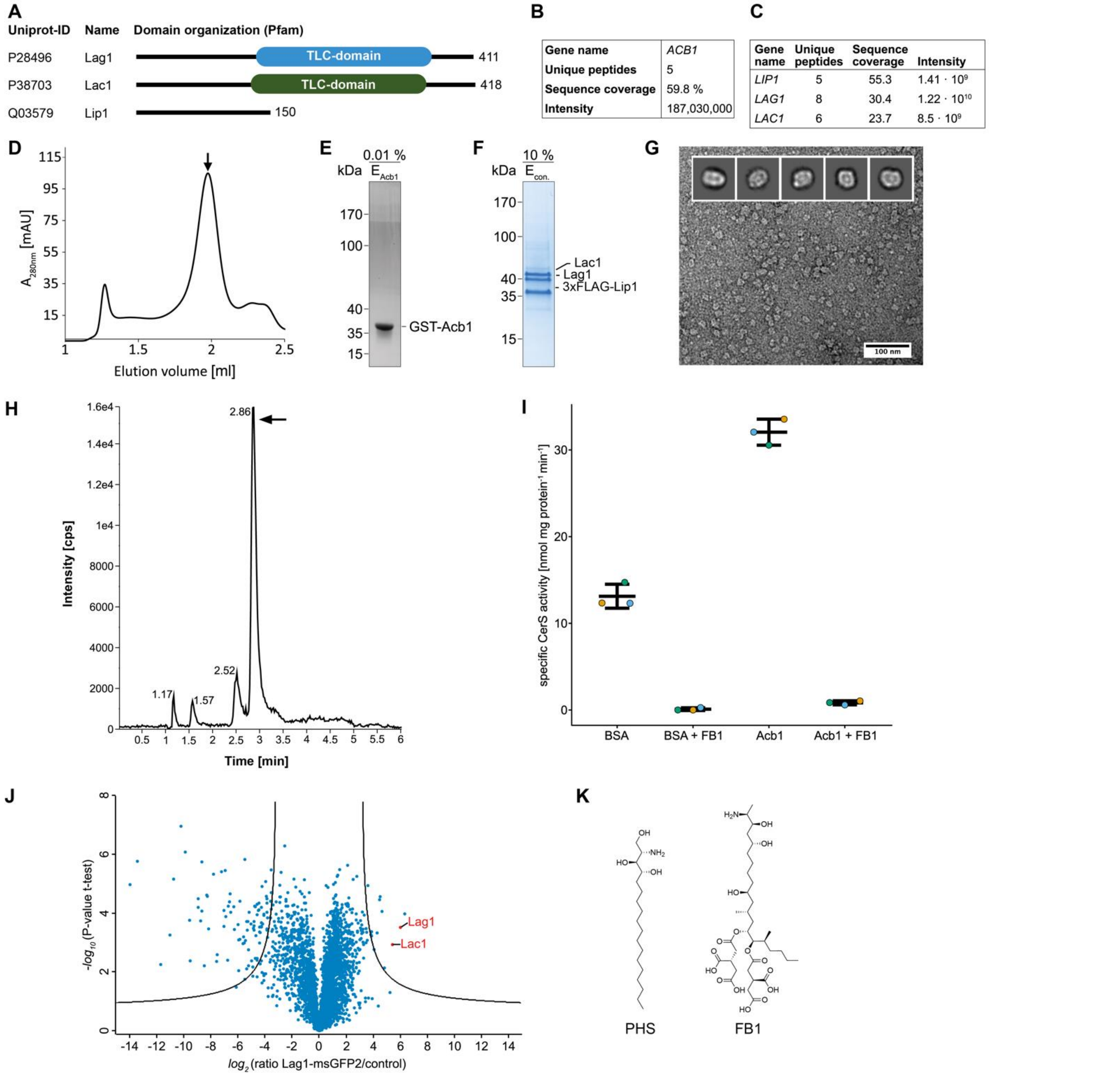


Sup-Fig. 1: ***In vitro* functional characterization of the yeast ceramide synthase complex**

**A** Pfam-based domain annotation of the CerS subunits. **B** Summary of the mass spectrometric analysis of the sample GST-Acb1 (E). **C** Summary of the mass spectrometric analysis of the CerS sample (F). **D** Size-exclusion chromatography profile of GDN-solubilized CerS complex. **E** Coomassie-blue stained SDS-PAGE gel of purified GST-Acb1. **F** Coomassie-blue stained SDS-PAGE gel of GDN-solubilized CerS complex of indicated fraction from (D). **G** Representative micrograph and 2D class averages from negative-stain TEM. 100 nm scale bar. **H** LC-MS/MS chromatogram of phytosphingosine 18:0 (PHS, m/z 318.499 → m/z 60) from purified CerS processed in the presence of the internal standard sphingosine d17:1 (m/z 286.274 → m/z 69). cps = counts per second. **I** Specific CerS enzyme activity measurement with or without the specific CerS-inhibitor fumonisin B1 (FB1). Control with addition of BSA and the Fatty-acid-Coenzyme A transport protein Acb1. n=3 technically independent samples for i and data are presented as mean values ± SD. **J** Label free proteomics of yeast cells expressing Lag1-msGFP2 compared to untagged control cells. In the volcano plot, the protein abundance ratios of Lag1-msGFP2 over control cells are plotted against the negative $log_{10}$ of the *P*-value of the two-tailed t-test for each protein. **K** Chemical structures of PHS and FB1.

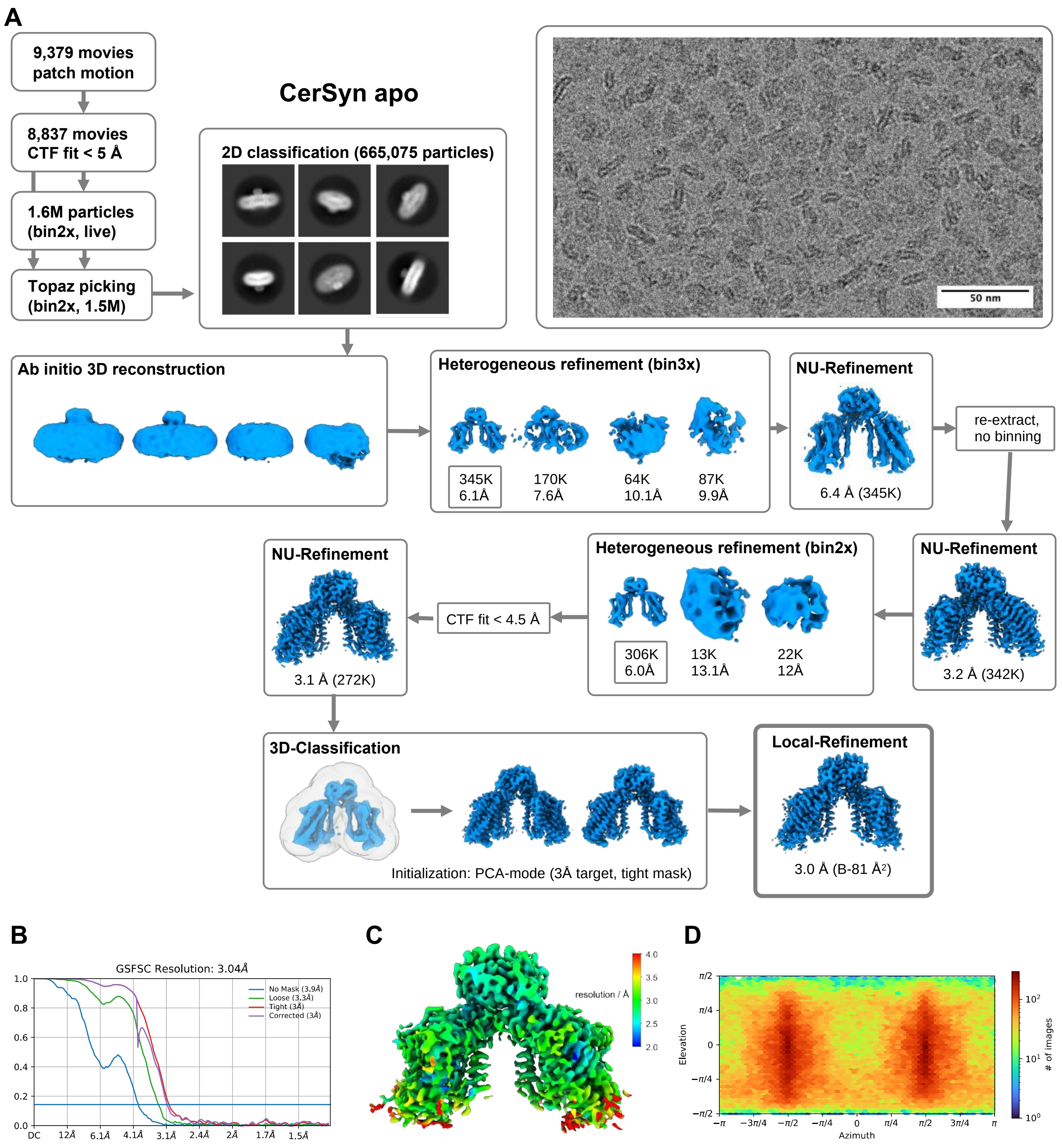


Sup-Fig. 2: **Cryo-EM analysis and validation of the ceramide synthase apo complex**

**A** Processing workflow for the CerS in its apo state. All processing steps were performed in cryoSPARC. Representative cryo-EM micrograph and 2D-class averages are given. 50 nm scale bar in micrograph. **B**

FSC-curve with gold-standard criterion at 0.134 and 3.0 Å overall resolution. **C** Local resolution estimation (threshold 0.16). **D** angular particle distribution.

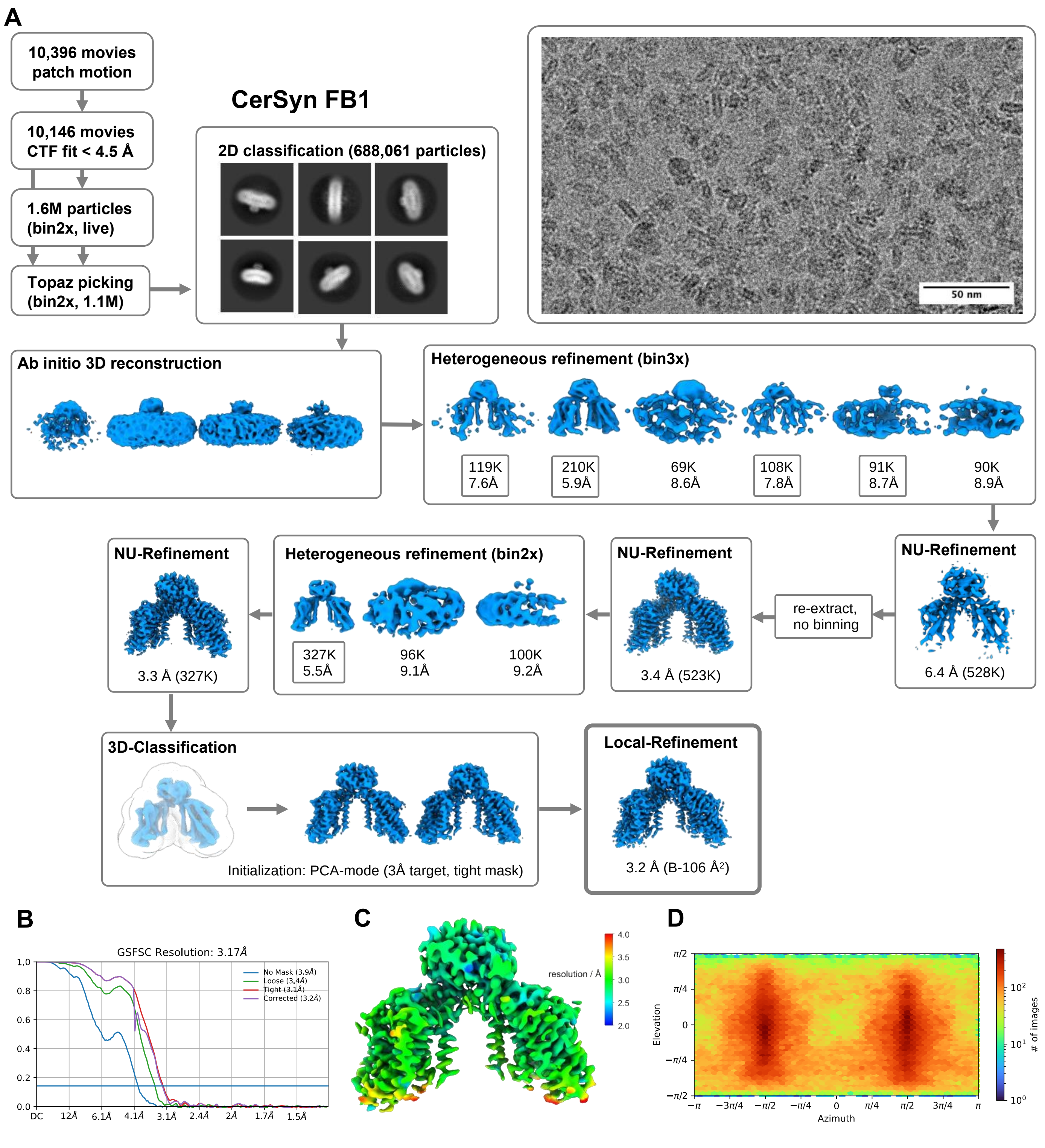


Sup-Fig. 3: **Cryo-EM analysis and validation of the FB1-bound ceramide synthase complex**

**A** Processing workflow for the FB1-bound CerS complex. All processing steps were performed in cryoSPARC. Representative cryo-EM micrograph and 2D-class averages are given. 50 nm scale bar in micrograph. **B** FSC-curve with gold-standard criterion at 0.134 and 3.2 Å overall resolution. **C** Local resolution estimation (threshold 0.16). **D** angular particle distribution.

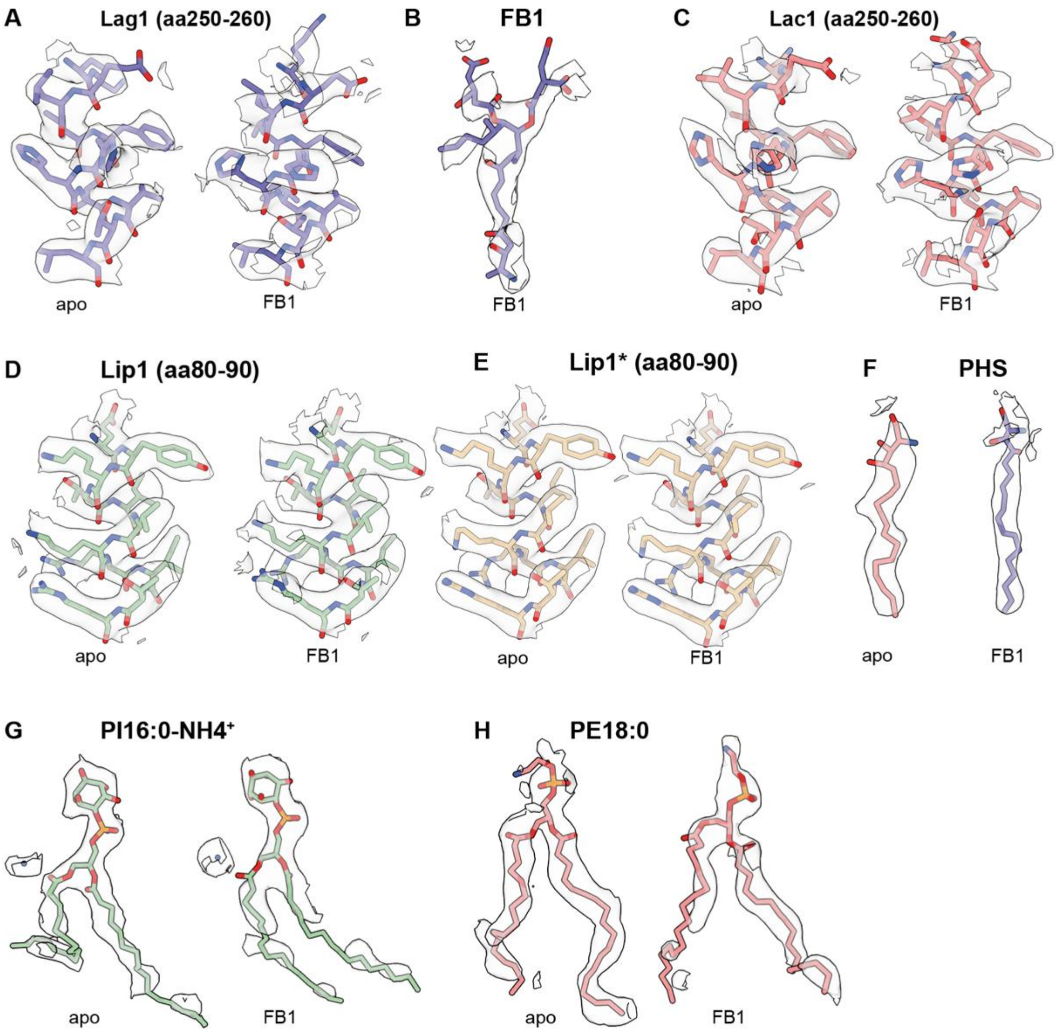


Sup-Fig. 4: **Local cryo-EM density map quality of apo and FB1-bound ceramide synthase**

Side-by-side comparison of selected residues and ligands within all cryo-EM density maps of **A** Lag1 **B** FB1 (fumonisin B1) **C** Lac1, **D** Lip1, **E** Lip1*, **F** PHS (phytosphingosine), **G** PI16:0-lipid (phosphoinositide) and ammonium ion $NH4^+$, **H** PE18:0-lipid (phosphoethanolamine). Contouring levels set to 0.16.

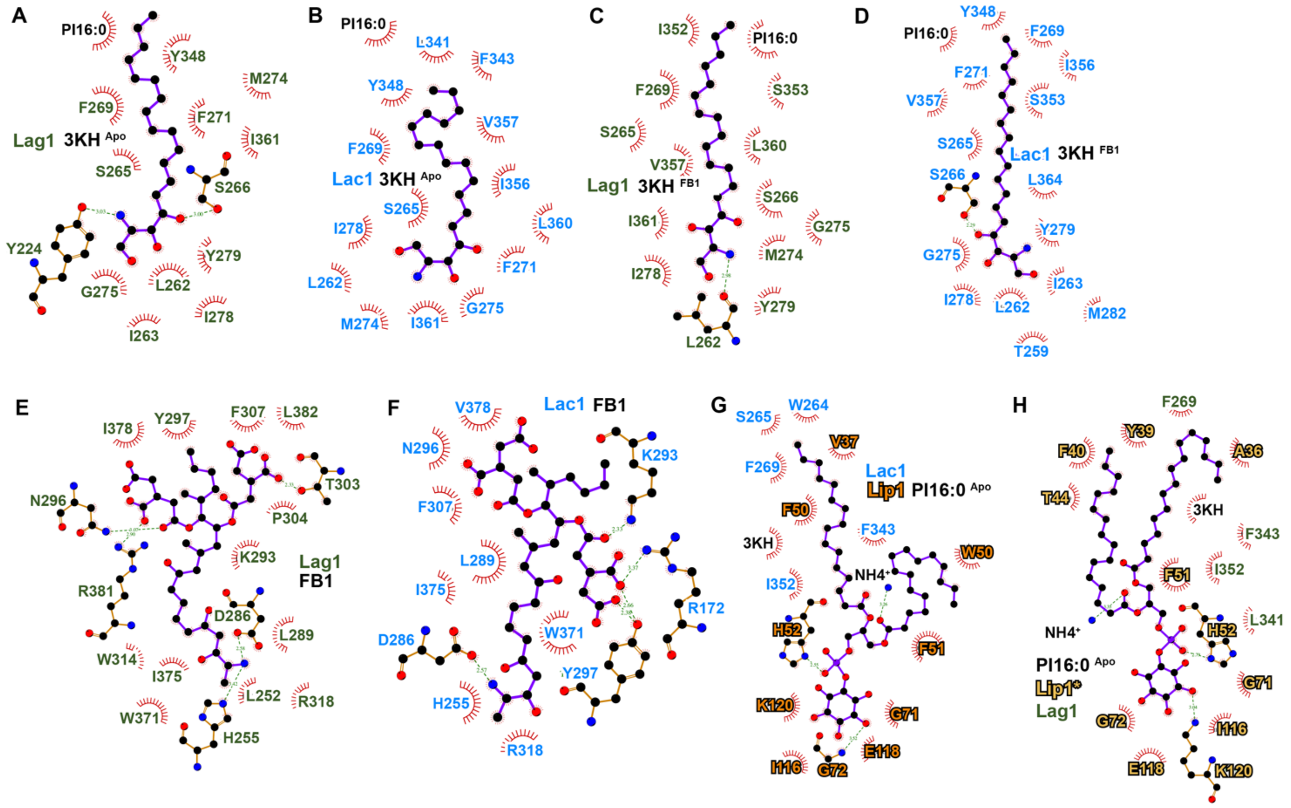


Sup-Fig. 5: **Ligand interaction diagram of ceramide synthase**

2D ligand interaction diagram of: **A** PHS in the apo-state with Lag1, **B** PHS in the apo-state with Lac1, **C** PHS in the FB1-bound state with Lag1, **D** PHS in the FB1-bound state with Lac1, **E** FB1-binding to Lag1, **F** FB1-binding to Lac1, **G** PI16:0 binding to Lac1 in the apo-state and **H** PI16:0 binding to Lac1 in the FB1-bound state. Diagrams were calculated and visualized with LigPlot+[1].

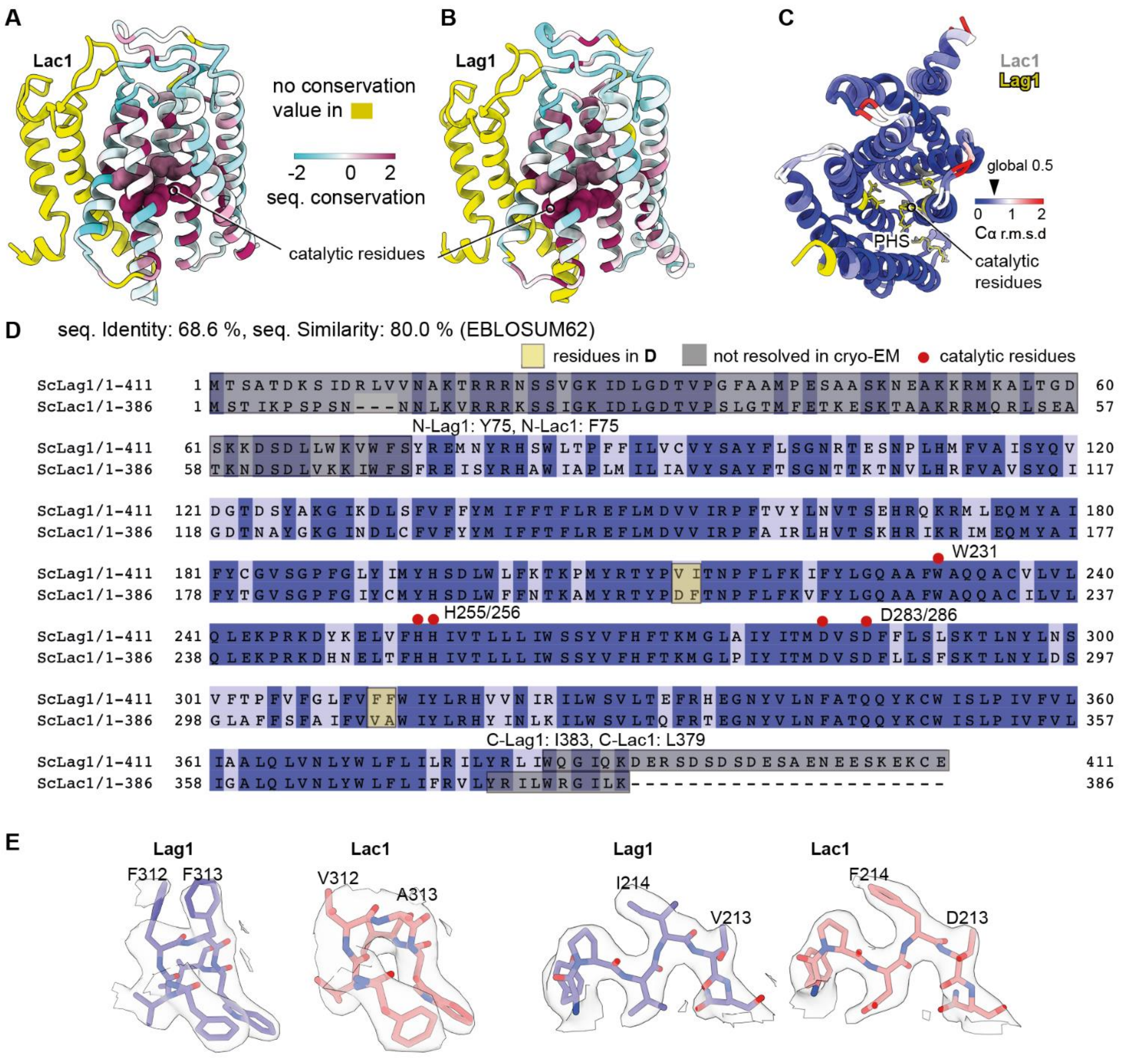


Sup-Fig. 6: **Structural conservation of the yeast ceramide synthase**

**A** Cartoon representation of Lac1 color-coded by sequence-conservation with data from Sup. Fig. 8A. **B** Cartoon representation of Lag1 color-coded by sequence-conservation with data from Sup. Fig. 9. Yellow areas denote regions of no conservation. The catalytic residues are highlighted as spheres. **C** Luminal view of the cartoon-representation of superimposed Lag1 (yellow) and Lag1 (grey), color-coded by Cα root-mean-square deviation (r.s.m.d.). Catalytic residues and PHS are shown as sticks. **D** Pair-based sequence alignment of Lac1 *Saccharomyces cerevisiae* (P28496) and Lag1 *Saccharomyces cerevisiae* (P38703). Sequences are color coded by conservation with a cut-off at 30 %. Catalytic residues are highlighted in red. Non-resolvable residues from cryo-EM are marked in light gray. Alignment was prepared with Jalview. **E** Comparison of local densities in Lag1 and Lac1 for density-based subunit differentiation. The corresponding sequences are highlighted in yellow in the alignment (D).

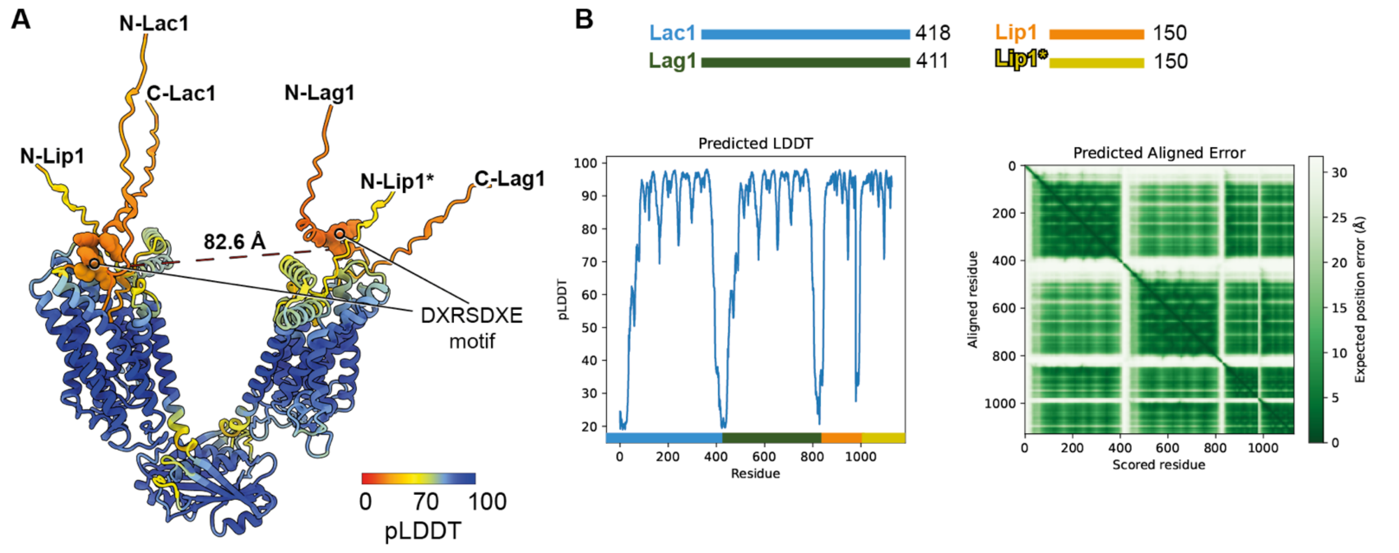


Sup-Fig. 7: **AlphaFold2 multimer prediction of the yeast ceramide synthase**

**A** AlphaFold2 multimer prediction of the yeast CerS (*Sc*Lag1: P38703, *Sc*Lac1: P28496, *Sc*Lip1: Q03579) shown as cartoon-representation, color-coded by the predicted local distance difference test (pLDDT). The low-confidence termini of each subunit are annotated accordingly. Distance measurement between Cα of Lag1-D390 and Lac1-D394 from their respective proposed dimerization motif (DXRSDXE). **B** Validation parameters of the prediction, including the pLDDT and the PAE (prediction aligned error matrix). The pLDDT plot is color-coded according to the different subunits. The prediction was run locally using AlphaFold v.2.1.0[2] in the multimer-mode (template date set to 2020-05-14).

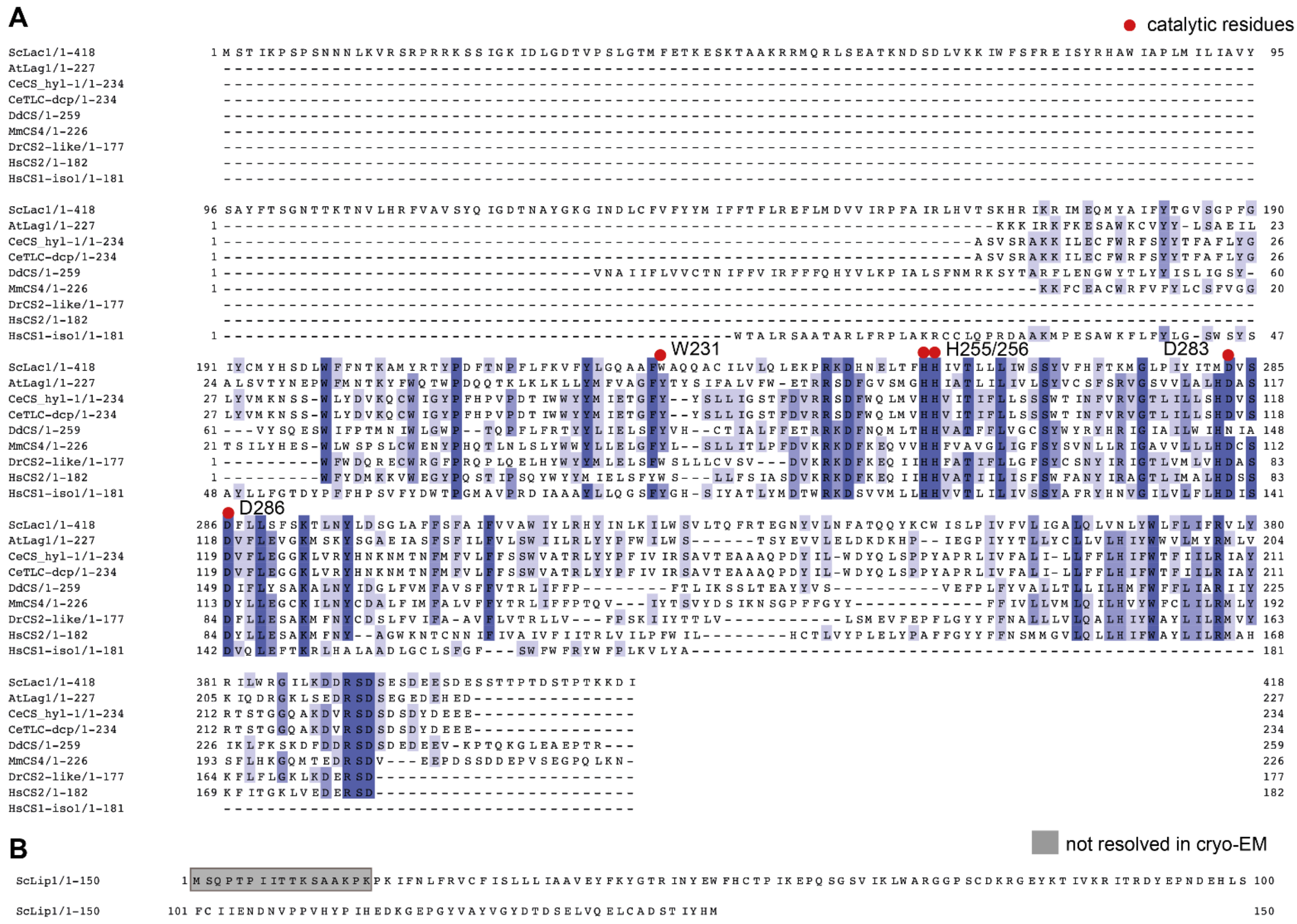


Sup-Fig. 8: **Sequence conservation of the yeast ceramide synthase subunits Lac1 and Lip1**

**A** Clustal-based multiple-sequence alignment (MSA) of Lac1 *Saccharomyces cerevisiae* (P28496), Lag1 *Arabidopsis thaliana* (Q9LDF2), CS hyl-1 *Caenorhabditis elegans* (G5ED45), TLC domain containing protein *Caenorhabditis elegans* (X5LPS5), CS *Dictyostelium discoideum* (Q54S87), CS4 *Mus musculus* (Q9D6J1), CS2-like *Danio rerio* (E7F9X7), CS2 *Homo sapiens* (Q96G23), CS1 *Homo sapiens* (P27544). Sequences are color coded by conservation with a cut-off at 30 %. Catalytic residues are highlighted in red. Sequences were selected by highest sequence identity hits from protein BLAST of selected model organisms. **B** Sequence of *Saccharomyces cerevisiae* Lip1 (Q03579). Non-resolvable residues from cryo-EM are marked in light gray. MSAs were prepared with Jalview[3].

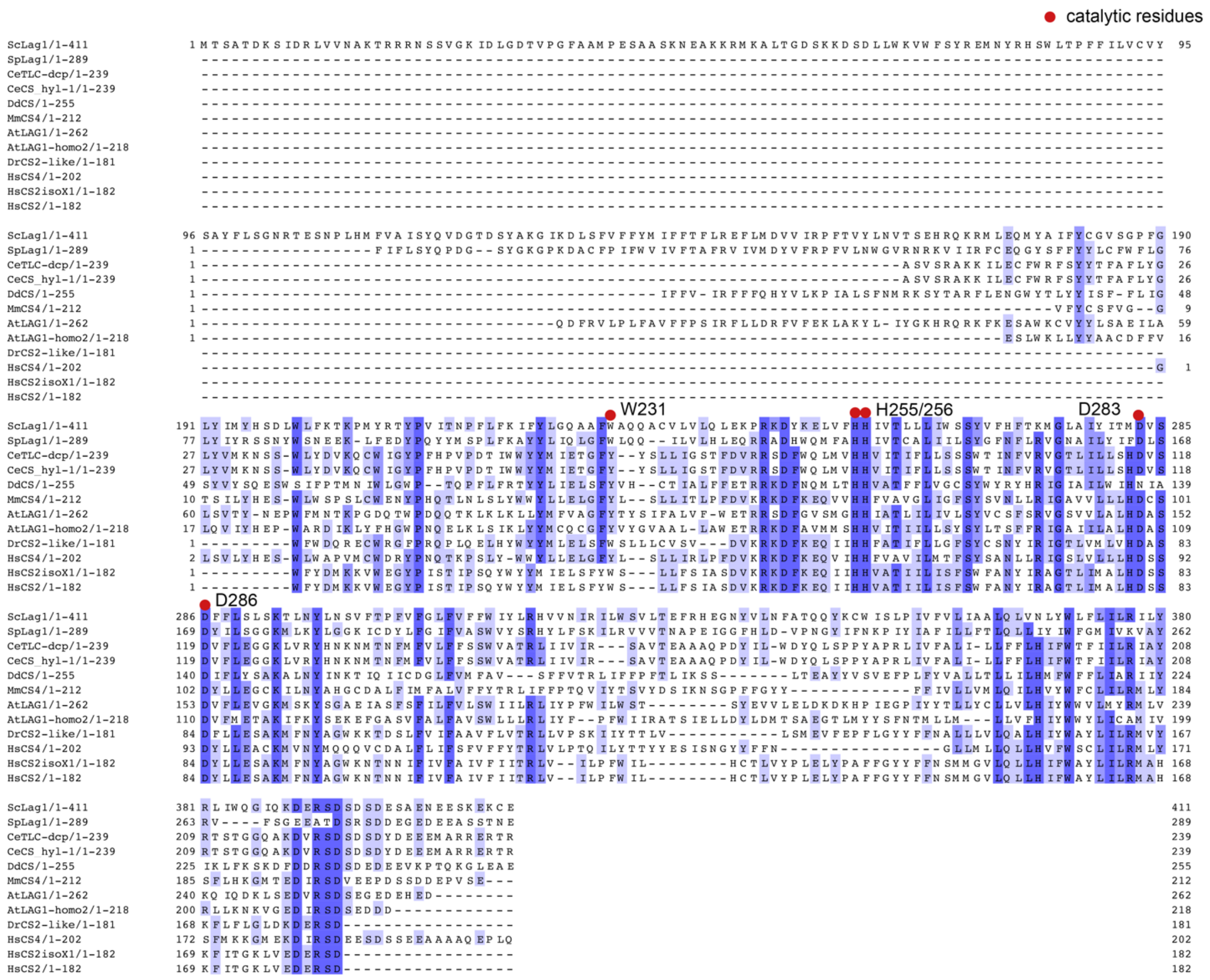


Sup-Fig. 9: **Sequence conservation of the yeast ceramide synthase subunit Lag1**

**A** Clustal-based multiple-sequence alignment (MSA) of Lag1 *Saccharomyces cerevisiae* (P38703), Lag1 *Saccharamyces pombe* (O13752), TLC domain containing protein *Caenorhabditis elegans* (X5LPS5), CS hyl-1 *Caenorhabditis elegans* (G5ED45), CS *Dictyostelium discoideum* (Q54S87), CS4 *Mus musculus* (Q9D6J1), Lag1 *Arabidopsis thaliana* (Q9LDF2), Lag1-homo2 *Arabidopsis thaliana* (A0A1I9LTU4), CS2-like *Danio rerio* (E7F9X7), CS4 *Homo sapiens* (Q9HA82), CS1 *Homo sapiens* (P27544), CS2 *Homo sapiens* (Q96G23). Sequences are color coded by conservation with a cut-off at 30 %. Catalytic residues are highlighted in red. Sequences were selected by highest sequence identity hits from protein BLAST of selected model organisms. MSAs were prepared with Jalview.

Sup-Tab. 1: **Cryo-EM data collection, refinement and validation statistics**

*Saccharomyces cerevisiae* ceramide synthase with Lag1, Lac1 and Lip1.

| | *Sc*CerS apo (PDB 8QTN, EMD-18652) | *Sc*CerS FB1 (PDB 8QTR, EMD-18653) |
|---|---|---|
| Data collection and processing | | |
| Microscope | TFS Glacios | TFS Glacios |
| Voltage (keV) | 200 | 200 |
| Camera | TFS Falcon 4i | TFS Falcon 4i |
| Energy filter | TFS Selectris | TFS Selectris |
| Slit width eV | 10 | 10 |
| Magnification (nominal) | 165,000 | 165,000 |
| Pixel size (Å/px) | 0.68 | 0.68 |
| Defocus range (µm) | 0.8-2.0 | 0.8-2.0 |
| Total exposure ($e^-$/Å$^2$) | 50 | 50 |
| Exposure rate ($e^-$/px/s) | 5.72 | 5.72 |
| Automation software | *EPU 2.9* | *EPU 2.9* |
| Processing software | *CryoSPARC* 4.3 | *CryoSPARC* 4.3 |
| Micrographs collected* | 9,379 | 9,379 |
| Micrographs used* | 7,967 | 7,967 |
| Final particle images | 159,440 | 259,645 |
| Point-group symmetry parameters | *C1* | *C1* |
| Resolution (global) (Å) FSC 0.134 | 3.0 | 3.2 |
| Map-sharpening *B* factor (Å$^2$) | – 81 | – 106 |
| Map-sharpening method | Global *B* factor | Global *B* factor |
| Refinement package | *phenix.real_space_refine* | *phenix.real_space_refine* |
| Model Composition | | |
| Non-H atoms | 15,520 | 15,582 |
| Protein Residues | 1,774 | 1,772 |
| Model refinement | | |
| Model-map scores | | |
| CC (mask) | 0.80 | 0.79 |
| CC (volume) | 0.76 | 0.76 |
| FSC model (0.5) | 3.0 | 3.5 |
| Av. grouped B factors (Å$^2$) | | |
| Protein residues | 38.71 | 62.34 |
| Ligands | 38.88 | 64.34 |
| R.m.s.d. from ideal values | | |
| Bond lenghths (Å) | 0.004 | 0.003 |
| Bond angles (°) | 0.521 | 0.591 |
| Validation | | |
| *MolProbity* score | 1.61 | 1.66 |
| CaBLAM outliers (%) | 1.74 | 1.75 |
| Clashscore | 10.04 | 9.44 |
| Poor rotamers (%) | 0.88 | 0.00 |
| $C^\beta$ outliers (%) | 0.00 | 0.00 |
| Ramachandran plot | | |
| Favored (%) | 97.59 | 97.11 |
| Allowed (%) | 2.41 | 2.89 |
| Outliers (%) | 0.00 | 0.00 |

* Single set of movies used for all reconstructions.

Sup-Tab. 2: **List of all yeast strains used in this study**

| Strain | Genotype | Reference |
|---|---|---|
| FF5102 | *MATa/MATα ura3-52 trp1Δ 2 leu2-3,112 his3-11 ade2-1 can1-100*; *GAL1-LAG1::natNT2; GAL1-LAC1::kanMX6; GAL1-3xFLAG-LIP1::URA3* | this study |
| FF5156 | *MATα leu2-3,112 ura3-52 his3-Δ200 trp- Δ901 lys2-801 suc2-Δ9 GAL* | this study |
| FF4201 | *MATα leu2-3,112 ura3-52 his3-Δ200 trp1-Δ901 ADE2 suc2-Δ9 GAL lys2-801 LAG1-msGFP2::natNT2* | this study |
| FF4196 | *MATα leu2-3,112 ura3-52 his3-Δ200 trp1-Δ901 ADE2 suc2-Δ9 GAL lys2-801 LAC1-msGFP2::natNT2* | this study |

Sup-Tab. 3: **List of all plasmids used in this study**

| Plasmid | Reference |
|---|---|
| pYM-N23 GAL1pr-3xFLAG::URA3 | Gonzalez-Montoro lab (Osnabrück) |
| pYM-N23 (GAL1pr)::natNT2 | Shvarev *et al*., (2022)[4] |
| pYM-N22 (GAL1pr)::kanMX4 | Shvarev *et al*., (2022)[4] |
| pYM msGFP2 NAT::natNT2 (5358) | Klössel *et al*., (2023) |

Sup-Tab 4: **List of all oligo nucleotides used in this study**

| Name | Sequence |
|---|---|
| S1 Lip1 | CTACAATTTTGCACATCAAAAAAGACCTCCAACTACCGAGTTTGTA AGTACAATGCGTACGCTGCAGGTCGAC |
| S4 Lip1 | GTT TTG GCT TAG CAG CTG ATT TTG TAG TTA TGA TGG GAG TGG GTT GAG ACA TCG ATG AAT TCT CTG TCG |
| S1 Lac1 | TTCATACCTCCGGTAAACATTTAGATAGACACAGTATCAATAAACA AGAGCTATGCGTACGCTGCAGGTCGAC |
| S2 Lac1 | GTGTAAAGAATTAATGTGTAATGGTTATACTACTTAAAAACACCGT TTTCCTTCAATCGATGAATTCGAGCTCG |
| S3 Lac1 | AAGTGATGAAAGTTCCACTACTCCAACTGACAGTACTCCAACGAA AAAGGATATTCGTACGCTGCAGGTCGAC |
| S4 Lac1 | CCT CAC TTT CAA ATT GTT GTT TGA AGG GCT TGG CTT TAT TGT CGA CAT CGA TGA ATT CTC TGT CG |
| S1 Lag1 | AGTGTTGAGAGTGAACTCCAAGATACAGAGAAACTGAAGAAATAA CGACAACATGCGTACGCTGCAGGTCGAC |
| S2 Lag1 | CTTCATACAGGGGGGAAATCATATGATGATACGTATTCTCCTTAA GATACGTTTAATCGATGAATTCGAGCTCG |
| S3 Lag1 | TGACAGTGATTCTGATGAGAGCGCTGAAAATGAAGAATCTAAGGA AAAGTGTGAACGTACGCTGCAGGTCGAC |
| S4 Lag1 | GTT TTT GCA TTA ACA ACT AAC CTA TCG ATA GAT TTG TCC GTA GCT GAT GTC ATC GAT GAA TTC TCT GTC G |

Sup-Tab. 5: **List of all exact *P*-values of measured ceramide synthase activities using One-way ANOVA with Turkey-test**

| **Measure** | ***P*-value** |
|---|---|
| Acb1+FB1 vs Acb1 | 0.000000002905748 |
| BSA vs Acb1 | 0.000000055701898 |
| BSA+FB1 vs Acb1 | 0.000000002424106 |
| BSA vs Acb1+FB1 | 0.000002246637493 |
| BSA+FB1 vs Acb1+FB1 | 0.817491290776557 |
| BSA+FB1 vs BSA | 0.000001437308980 |

**Sup-Tab. 6:** List of used transitions for targeted lipidomics

| **Description** | **Q1 mass** | **Q2 mass** | **CE (V)** | **CXP (V)** | **DP (V)** | **EP (V)** |
|---|---|---|---|---|---|---|
| Sphingosine d17:1 | 286.274 | 69 | 55 | 10 | 51 | 10 |
| Phytosphingosine 18:0 | 318.300 | 60 | 45 | 10 | 166 | 10 |
